# From coral atolls to coal spoil heaps: two new species and two dark fungal taxa expand the monospecific genus *Innospora* (*Paraglomerales*, *Glomeromycota*)

**DOI:** 10.64898/2026.11.16.766956

**Authors:** Franco Magurno, Sylwia Szczecińska, Demetra Di Loreto, Dewi Fatmawati, Keyvan Esmaeilzadeh-Salestani, Maurice Wong, Mathieu Favoreu, Leho Tedersoo, Thomas Crossay

## Abstract

Morphological and phylogenetic analyses based on partial rDNA sequences supported the description of two new species of *Innospora*: *Innospora atollensis* sp. nov. and *Innospora fodinalis* sp. nov. These species expand the monospecific genus *Innospora,* which was formerly represented only by *I. majewskii*. *Innospora atollensis* was discovered on a coral atoll in French Polynesia, whereas *I. fodinalis* was recovered from a post-coal-mining spoil heap in Poland. Spores of *I. atollensis* and *I. fodinalis* are colorless, small and have a spore wall with three layers as in *I. majewskii*. Morphologically, the spores of *I. fodinalis* differ from those of other *Innospora* species by the presence of granular ornamentation on the first layer of the spore wall, whereas those of *I. atollensis* differ by having the third layer of the spore wall significantly thicker, and by the presence of a closed septum. Analysis including environmental sequences extended the known distributions of both species to several additional sites, with *I. atollensis* recorded in the Caribbean and Australia and *I. fodinalis* across the Eurasian continent. In addition, we detected two eDNA lineages at the level of species that meet the International Standard to Catalogue Dark Fungal Taxa criteria for designation as dark fungal taxa. These are formally described as *Innospora austroafricana*-ENAS and *Innospora palaearctica*-ENAS. The former has been recorded exclusively in South Africa, whereas the latter exhibits a broader distribution across the Euro-Mediterranean region.

## Introduction

The order *Paraglomerales* (class *Paraglomeromycetes* within the phylum *Glomeromycota*) currently comprises two families, *Paraglomeraceae* and *Pervetustaceae*, and three genera: *Paraglomus*, *Innospora*, and *Pervetustus*. Before the introduction of molecular markers, members of *Paraglomerales*, like most glomeromycotan fungi producing glomoid spores, were broadly classified within the genus *Glomus*. Morton and Redecker (2001), combining morphological characteristics of *G. occultum* and *G. brasilianum* with rDNA sequence data, fatty acid profiles, and immunological reactions, established the genus *Paraglomus* and the family *Paraglomeraceae*. Members of this lineage have been described as characterised by small, pale, glomoid spores, with one or two walls, each wall comprising two to three layers; the spores are generally produced singly at the tip of subtending hypha (Morton and Redecker 2001; de Mello et al. 2013; Błaszkowski et al. 2017). Concurrently, Schüßler et al. (2001) recognized *Paraglomerales* as a distinct order in the newly erected phylum *Glomeromycota*. Subsequently, Oehl et al. (2011) established the class *Paraglomeromycetes* to accommodate *Paraglomerales* based on phylogenetic evidence. Although *Paraglomeromycetes* are considered the earliest-diverging lineages within *Glomeromycota*, recent phylogenetic studies have not consistently supported this placement (Wijayawardene et al. 2025).

Molecular studies revealed that *Paraglomus majewskii*, a species described by Błaszkowski et al. (2012), was genetically highly divergent from other *Paraglomus* species and formed a distinct lineage within *Paraglomerales*, despite the morphological similarity of the hyaline, smooth spores and the simple wall structure. These findings prompted the taxonomic revision by Błaszkowski et al. (2017), in which *Pa. majewskii* was transferred to the newly established monospecific genus *Innospora*, resulting in the new combination *I. majewskii*. A clear morphological characteristic distinguishing *I. majewskii* from other *Paraglomus* species is the presence of a single-walled spore; however, this trait is not unique to *Innospora*, as single-walled spores also occur in other glomoid genera (de Mello et al. 2013; Błaszkowski et al. 2017). Because of the morphological simplicity of their spores, the diversity of *Paraglomerales* may be underestimated in spore-based surveys. In eDNA surveys, underestimation may result from poor amplification of the divergent rDNA of some *Paraglomerales* lineages by *Glomeromycota*-specific primer pairs widely used in metabarcoding, such as NS31/AM1 (Helgason et al. 1998). Nevertheless, long-read eDNA datasets generated with broader fungal or eukaryotic primers, such as those compiled in EUKARYOME (Tedersoo et al. 2024), contain *Paraglomerales* lineages that are not known from cultures or spores. Their formal recognition, however, remains constrained by nomenclatural requirements.

Under the International Code of Nomenclature for algae, fungi, and plants (Turland 2025), a name can be validly published only with a physical type. Proposals to allow DNA sequences to serve as types were rejected at the International Mycological Congresses in 2018 and 2024 (Lücking et al. 2021). Fungi known only from eDNA, termed dark fungal taxa (DFT), therefore cannot be named under the Code. In *Glomeromycota*, environmental lineages have mostly been communicated as virtual taxa (Öpik et al. 2014) or as UNITE species hypotheses (SH; Kõljalg et al. 2013). Their identifiers are stable but awkward for human communication and are not connected to Linnaean names. The International Standard to Catalogue Dark Fungal Taxa (ISCDFT; Aime et al. 2026) provides an alternative outside the Code. Under it, a DFT receives a Linnaean-style designation with the suffix “-ENAS” (environmental nucleic acid sequence). Each designation is fixed to a single reference sequence linked to a persistent UNITE SH DOI, diagnosed by positional nucleotide differences, supported by phylogenetic evidence from several independent samples, and registered in a nomenclatural repository. A designation can later be validated under the Code once type material becomes available, for example from trap cultures established with soil from sites where the DFT occurs.

In the present study, we describe two new species of *Innospora*, *I. atollensis* sp. nov. from a coral atoll in French Polynesia, and *I. fodinalis* sp. nov. from a post-coal-mining spoil heap in Poland. Both species descriptions are based on the morphology of spores from single-species cultures and phylogenetic analyses of partial SSU–ITS–LSU sequences. In addition, we analysed environmental sequences from public databases with two aims: (i) to assess the geographic distribution and habitat range of the newly described *Innospora* species, and (ii) to detect further species-level eDNA lineages that could meet the ISCDFT criteria to be designated as dark fungal taxa.

## Material and methods

### Study sites

The Rangiroa atolls is located in the north-western part of the Tuamotu archipelago in French Polynesia, in the central South Pacific (14°58’S; 147°37’W, Fig. 1A). This atoll consists of a narrow strip of land surrounding a central lagoon. The land edge is segmented by shallow channels, known locally as ‘hoa’, which lead to islets known as ‘motu’ (Fig. 1B). Most of the ‘hoa’ are shallow passages linking the ocean and the lagoon. Along the edges of these atolls are numerous isolated shallow ponds, most of which are less than a meter deep and vary in size from a few dozen to several hundred square meters. These ponds are the result of the combined action of erosion and sedimentation (Duvat et al. 2020, 2022).

**Figure 1.**
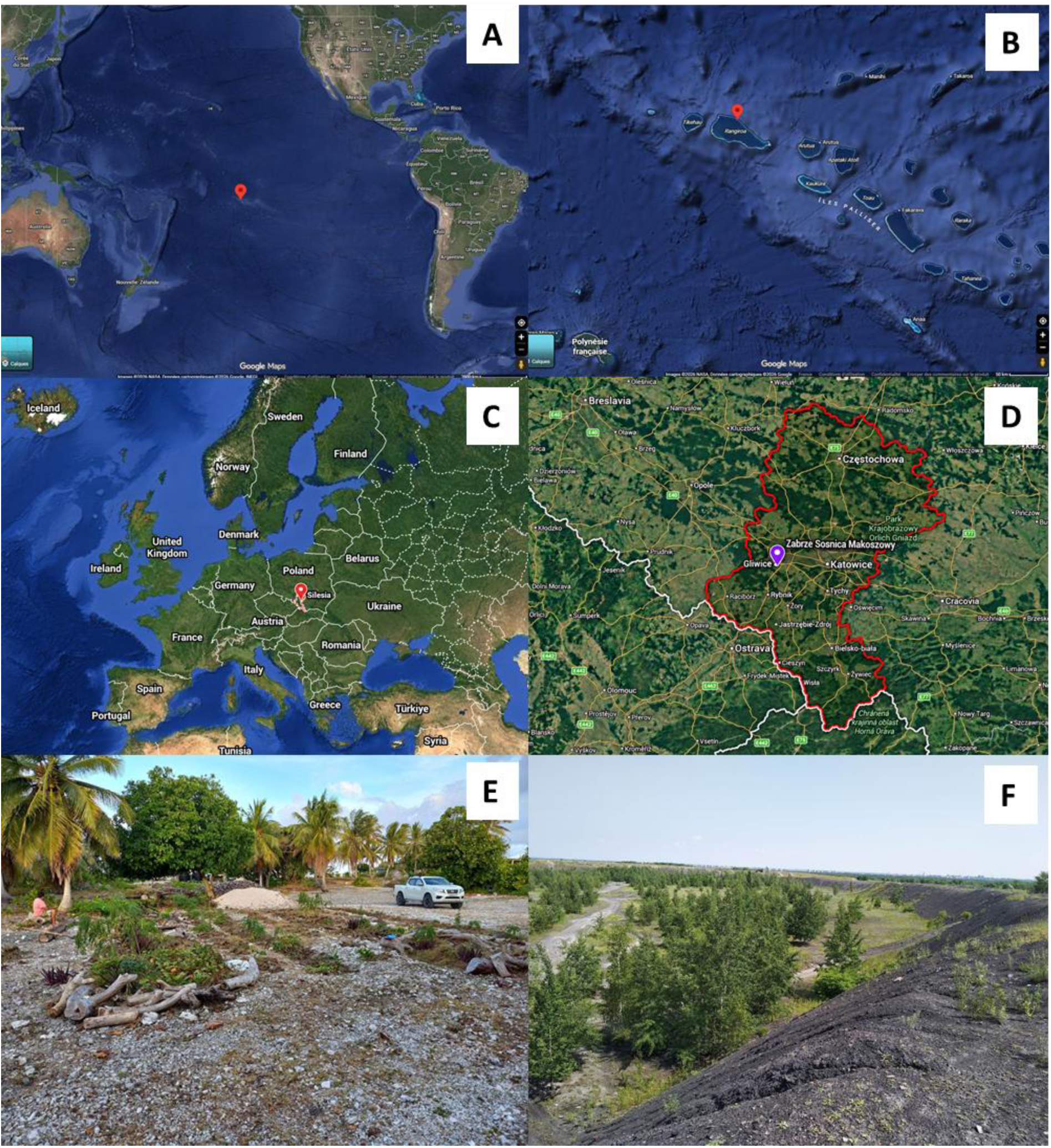
**A** Location of French Polynesia. **B** Location of the atoll of Rangiroa. **C** Silesian Voivodship, Poland. **D** Zabrze Sosnica Makoszowy site in Silesia, Poland. **E** Rangiroa sampling site. **F** Zabrze Sosnica Makoszowy spoil heap sampling site.

Rangiroa is the second largest atoll in the world, with an area of 1,763 km^2^. The land area of Rangiroa (66 km^2^) is made up of islets of varying sizes. Low-lying islets have soil composed mainly of pebbles and sand, and some are even completely devoid of vegetation or covered with low, often ephemeral, sometimes shrubby plants. The largest islets are generally surmounted by basal platforms of coral conglomerate, up to 1 to 4 m thick, generally colonized by dense, shrubby to tall vegetation, including remnants of native plants. These islets are known as wooded islets. Relatively numerous on Rangiroa, wooded islets are apparently typical of Tuamotu atoll lagoons compared with other Pacific atolls. In Rangiroa, the air temperature ranges from 22°C to 31°C. Rainfall is around 1,500 mm/year, with values ranging from 900 to 2,200 mm (Duvat et al. 2020, 2022).

In the study site (Fig. 1E), the vegetation is a mosaic of shrubby autochthonous vegetation and agronomic species. There are several unifying features: shrubby vegetation is evergreen, 1–7 m tall, more or less bushy, most species are sclerophyllous. The vegetation is dominated by *Cocos nucifera* L., 1753, *Ficus carica* L. and *Carica papaya* L. Most of the plant species have a well-developed superficial root system to grow on shallow soil composed mainly of coral debris and sand (Fig. 1E).

The Upper Silesian Coal Basin is located primarily in the Silesian Voivodeship of southern Poland (Fig. 1C, D) and is often referred to as “Black Silesia” because of its long history of mining, particularly of hard coal, zinc, and lead ores. The mining landscape has been transformed by the extraction of billions of tons of hard coal, waste rock and sand, which was used for hydraulic backfilling of underground workings (Dulias 2024). The region is characterized by a temperate climate, with a mean annual precipitation of 727 mm and an average annual temperature of 8.5 °C (Pietrzykowski et al. 2026). One of the most visible consequences of mining is the formation of spoil heaps composed of overburdened material and heterogeneous mining waste. Historically, these post-mining spoil heaps were left to undergo spontaneous revegetation and therefore represent a characteristic landmark of the Silesian landscape. Spoil heap soils are often characterized by an unfavorable granulometric structure, consisting predominantly of bedrock fragments and gravel, high bulk density, low pH, limited nutrient availability, high salinity, low water-holding capacity, and low biomass productivity (Woźniak 2010).

The sampling area was located on the Zabrze Sosnica Makoszowy spoil heap (Fig. 1F), an active post-coal-mining wasteland covering approximately 170 hectares, located at an altitude of over 310 m above sea level (50°16′22” N; 18°44′43” E). On the heap, spontaneous revegetation ranges from the early stages of primary succession to intermediate successional stages, depending on when the mining waste was deposited. The vegetation is represented by a mosaic of communities occurring in patches and dominated by herbaceous plant species such as *Calamagrostis epigejos* (L.) Roth, 1788, *Arrhenatherum elatius* (L.) P.Beauv. ex J.Presl & C.Presl, 1819, *Centaurea stoebe* Ledeb., 1833, *Chamaenerion palustre* Schreb., 1771, *Festuca arundinacea* Schreb., 1771, *F. rubra* L., 1753, *Medicago lupulina* L., 1753, *M. sativa* L., 1753, *Melilotus albus* Medik., 1787, *Phragmites australis* (Cav.) Steud., 1840, *Tussilago farfara* L., 1753; and tree species such as *Betula pendula* Roth, 1788, and *Populus tremula* L., 1753 (Bierza et al. 2023).

### Sampling, establishment, and growth of single-species cultures

Spores of the French Polynesian morphotype were sampled and used to establish single-species cultures according to the method described by Crossay et al. (2018). Spores were isolated from the rhizospheric soil of *Ficus carica* L. sampled on Rangiroa Island. Then, single-species pot cultures were prepared by inoculating a substrate consisting of a 3-times autoclaved mix of peat moss (30%), river sand (35%), vermiculite (30%), and bentonite (5%). A pot was then inoculated with 100 spores of the same morphotype, using *Sorghum vulgare* as the host plant. Eight months after spore inoculation, several spore extractions and microscopic examinations were performed to confirm that only one morphotype was present in the culture pot.

Rhizospheric soil of *Tanacetum vulgare* collected at Zabrze Sosnica Makoszowy site (Poland) was used for inoculation of pot trap culture with *Plantago lanceolata* L. as host plant, and autoclaved quartz sand amended with 5% bentonite clay as substrate. After approximately five months, spores were extracted from the trap culture and used to establish pure-species pot cultures inoculated with single spores, using plant host and substrate as above. Pot cultures were maintained in a plant growth room at the Institute of Biology, Biotechnology and Environmental Protection (Katowice, Poland) and re-established approximately every six months upon verification of the presence of spores.

### Morphological analyses

Spores from single-species cultures were extracted as previously described in Crossay et al. 2018 using the methods of Daniels and Skipper 1982. Morphological features of spores, and phenotypic and histochemical characters of spore wall layers were determined after examination of at least 100 spores mounted in water, lactic acid, PVLG (Omar et al. 1979), and a mixture of PVLG and Melzer’s reagent (1:1, v/v). The terminology for spore subcellular structures follows Błaszkowski (2012). Spore color was examined under a dissecting microscope on fresh specimens immersed in water. Colors were determined using the INVAM color chart. Photographs were taken using a digital camera (Leica DFC 295) on a compound microscope (Olympus BX 50) equipped with *Nomarski* differential interference contrast optics and Leica Application Suite Version V 4.1 software. Specimens of Rangiroa and Polish isolates, mounted in PVLG and in a (1:1) mixture of PVLG and Melzer’s reagent, were deposited at the Mycological Herbarium of the National Museum (France, Paris) and in the KTU Herbarium of the University of Silesia (Poland, Katowice), respectively. All morphological comparisons of *Innospora* were performed using species descriptions of Błaszkowski et al. (2012) and Błaszkowski et al. (2017).

### Molecular analyses

Total genomic DNA extraction was performed from ca. 100 spores per isolate with DNeasy PowerSoil Pro Kit (Qiagen, Hilden, Germany) as described in Magurno et al. (2024). Both primers described in Krüger et al. (2009) and newly designed primers were tested to obtain partial SSU-ITS-LSU rDNA (hereafter 45S) sequences from the isolates. Two reverse primers (Silesia_Rev: GTT TAA AGC CAT TAC GTC AAC G; Rangiroa_Rev: CCG CAA CCT GGT GTT CAA AG) were designed according to the following strategy. First, a partial LSU fragment was amplified using the Ful-F and Ful-R primers (Malicka et al. 2022). Then, the sequences were used to query the nucleotide sequence repository EUKARYOME v.2.1 (Tedersoo et al. 2024) and retrieve matches with a high percentage of identity. The primers were designed in the hypervariable D2 region in the proximity to the FUL-R site, based on consensus regions in the alignment between isolates and EUKARYOME sequences. Primers were assessed using the OligoAnalyzer™ tool (Integrated DNA Technologies, IDT) for temperature of annealing, hairpins and dimers. Primer specificity was verified using NCBI Primer-BLAST vs non-*Glomeromycota* sequences. The two reverse primers were used in combination with the universal fungal primer V9G (TTA CGT CCC TGC CCT TTG TA, de Hoog and Gerrits van den Ende 1998) with site of annealing immediately downstream of SSUmCfx (Krüger et al. 2009). Similarly, in order to amplify part of the RPB1 gene, the primers RPB1_innospora_forw (CGT CCA AGT ATA CAA ATG GA) and RPB1_innospora_rev (ACT CTG TGA CCC ATC ATT GAC A) were designed on the exon 4 and 5, respectively, from conserved regions identified in an alignment comprising all available *Paraglomerales* RPB1 sequences and the RPB1 sequence extracted from the *Innospora majewskii* genome (DOI: 10.5281/zenodo.13963845).

All PCR reactions were performed using the Phusion Plus DNA Polymerase (Thermo Fisher Scientific, Waltham, MA, USA) with a universal annealing temperature of 60 °C, according to the producer’s instructions. Purification of PCR products, cloning, screening, plasmid extraction and sequencing were as in Crossay et al. (2026). Sequences were deposited in GenBank with the accession numbers: (45S: PZ835639 - PZ835643, PZ804334 - PZ804335, PZ738842 - PZ738845; RPB1: QB246059).

### Phylogenetic analyses

Phylogenetic analyses were conducted by assembling two sequence datasets. The first dataset included the two putative new species, the nine known species with available sequences for the three genera recognized in the *Paraglomerales* (*Innospora*, *Pervetustus*, *Paraglomus*) and, as outgroup, species representing the four established orders in the class *Archaeosporomycetes* (Oehl et al. 2026). The second dataset included members of *Paraglomeraceae* and environmental sequences retrieved with blastn from GenBank and EUKARYOME v.2.1 databases. The additional environmental sequences were retrieved by exploring the 90% molecular identity neighbourhood of the two new *Innospora* species, potentially encompassing conspecific sequences and neighbouring lineages. Overall, the datasets comprised 47 and 90 sequences, respectively. Sequences were aligned using MAFFT v.7 (Katoh and Standley 2013) using the E-INS-i iterative refinement algorithm. Then, alignments were inspected and edited in MEGA v5 (Tamura et al. 2011) to remove overhanging ends and rule out poorly aligned regions. Both datasets were used for maximum likelihood (ML) and Bayesian inference (BI), following the partitioning strategy and substitution models described by Esmaeilzadeh-Salestani et al. (2025).

Maximum-likelihood analyses were conducted using RAxML-NG (Kozlov et al. 2019) with branch support estimated from 1,000 bootstrap replicates. Bayesian phylogenetic inference was performed in MrBayes v.3.2.6 (Ronquist et al. 2012), with two independent Markov chain Monte Carlo (MCMC) runs, each with four chains for 10 million generations, a stop rule of standard deviation of split frequencies = 0.005/0.009, and sampling every 1,000 generations.

Phylogenetic trees were visualized in FigTree v.1.4.4 (Rambaut 2018). Clades were considered well supported when Bayesian posterior probabilities were ≥ 0.95 and maximum-likelihood bootstrap values were ≥ 70%.

### Diagnostic barcodes

Diagnostic barcodes for the two new species and the two DFT were identified in the ITS2 and LSU subregions of the second alignment. A barcode was defined as a short oligonucleotide motif (∼20 bp) present in all sequences of the target lineage and differing by at least two nucleotides from the homologous region of all other *Innospora* sequences. Barcode positions are given relative to a representative 45S sequence of each isolate (new species) or to the reference sequence (DFT). The specificity of each barcode was verified by searching it against EUKARYOME v.2.1 and GenBank, and only motifs found exclusively in sequences of the target lineage were retained.

### Designation of dark fungal taxa

Environmental lineages were designated as dark fungal taxa (DFT) following the International Standard to Catalogue Dark Fungal Taxa (ISCDFT; Aime et al. 2026). A lineage was considered a candidate DFT when it formed a well-supported clade distinct from all described species and was represented by at least five sequences from at least three independent samples. Mean intraspecific identity and identity to the closest neighbouring lineage were calculated from global pairwise alignments of all 45S sequences in the second dataset. For each DFT, the longest sequence with the fewest deviating base calls within the lineage was selected as the reference sequence. Diagnoses are based on the diagnostic barcodes described above, which are present in the reference sequence. Reference sequences will be registered in the UNITE Species Hypotheses database (Kõljalg et al. 2013; Abarenkov et al. 2024) during manuscript revision, and the resulting SH DOIs will be used to register the designations in MycoBank.

## Results

### Molecular data and phylogenetic analysis

Several PCR attempts were unsuccessfully conducted with the primer set of Krüger et al. (2009). Conversely, partial 45S amplicons were obtained from the two isolates using the newly designed reverse primers in combination with the forward primer V9G.

Blastn analysis indicated that the isolates from Rangiroa atoll in French Polynesia and from Zabrze Sosnica Makoszowy spoil heap (Poland) represented two potential new species of *Innospora.* The intra-isolate diversity, calculated from the sequences obtained for each isolate, was low, within a range of 0.25-0.30%.

The Polynesian and Polish isolates differ from *Innospora majewskii* by percentage dissimilarities of ca. 20% and 14%, respectively, and by ca. 19% between each other (45S). Considering the LSU alone, the percentage dissimilarity with *Innospora majewskii* drops to ca. 8% and 6%, respectively, and 8% between each other, values comparable to the distances among *Paraglomus* species.

A partial RPB1 sequence was obtained only for the Polish isolate, and therefore, this molecular marker was not employed in the phylogenetic analysis. This sequence shared ca. 2.5% dissimilarity with the sequence of *I. majewskii*, whereas the values from the comparisons between the sequences available for three *Paraglomus* species (*Pa. brasilianum*, *Pa. occultum*, *Pa. laccatum*) ranged from ca. 4.2% to 9.9%.

Maximum likelihood (ML) and Bayesian inference (BI) phylogenies of the dataset including species of *Paraglomerales* recovered a congruent tree topology, with the two isolates representing independent, fully or highly supported lineages within the *Innospora* genus (Fig. 2). The two *Innospora* species, hereafter referred to as *I. atollensis* (from Rangiroa atoll) and *I. fodinalis* (from Poland), and *I. majewskii* formed a highly supported clade (ML = 98%, BI = 1.00) without any supported sister relationships observed among the three species.

**Figure 2.**
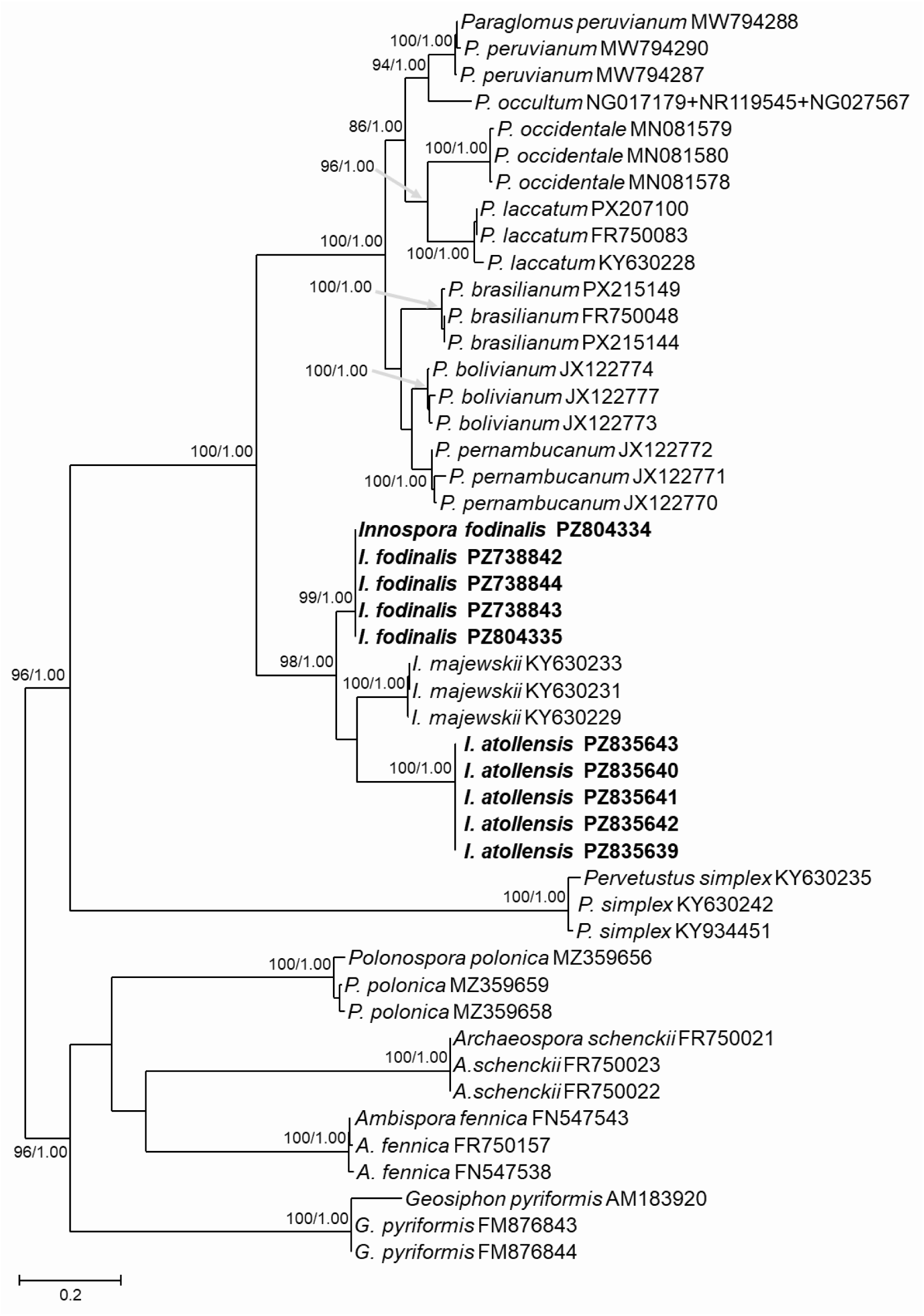
Consensus tree showing the phylogenetic placement of the two new *Innospora* species (indicated in bold) among members of *Paraglomerales*. For the Bayesian inference, all convergence diagnostics indicated that convergence had been achieved (ASDSF = 0.005; average PSRF for parameter values = 1.000). Values on the branches represent maximum-likelihood bootstrap support (BS) and Bayesian posterior probabilities (PP); only BS ≥ 85% and PP ≥ 0.97 are shown. The scale bar indicated 0.2 substitutions per site.

A congruent tree topology was also obtained in the maximum-likelihood (ML) and Bayesian inference (BI) phylogenies of the second dataset, including the three *Innospora* species, 58 environmental sequences recovered by blastn queries, and *Paraglomus* species as the outgroup (Fig. 3). Environmental sequences either grouped with the three species clades or formed independent lineages. All species clades enlarged with environmental sequences retained full or high support. Among them, *I. atollensis* included the highest number of environmental sequences, reflecting the highest intra-clade variability (% dissimilarity = ca. 4%), and a trans-Pacific–Caribbean distribution with several occurrences in the Caribbean, Australia and French Polynesia, as shown in Suppl. material 1. *Innospora fodinalis* showed a slightly lower intra-clade variability (% dissimilarity = ca. 3.7%) and occurrences across Eurasia, including two post-mining wasteland sites (in Southern Poland and France) such as the sampling site of the isolate.

**Figure 3.**
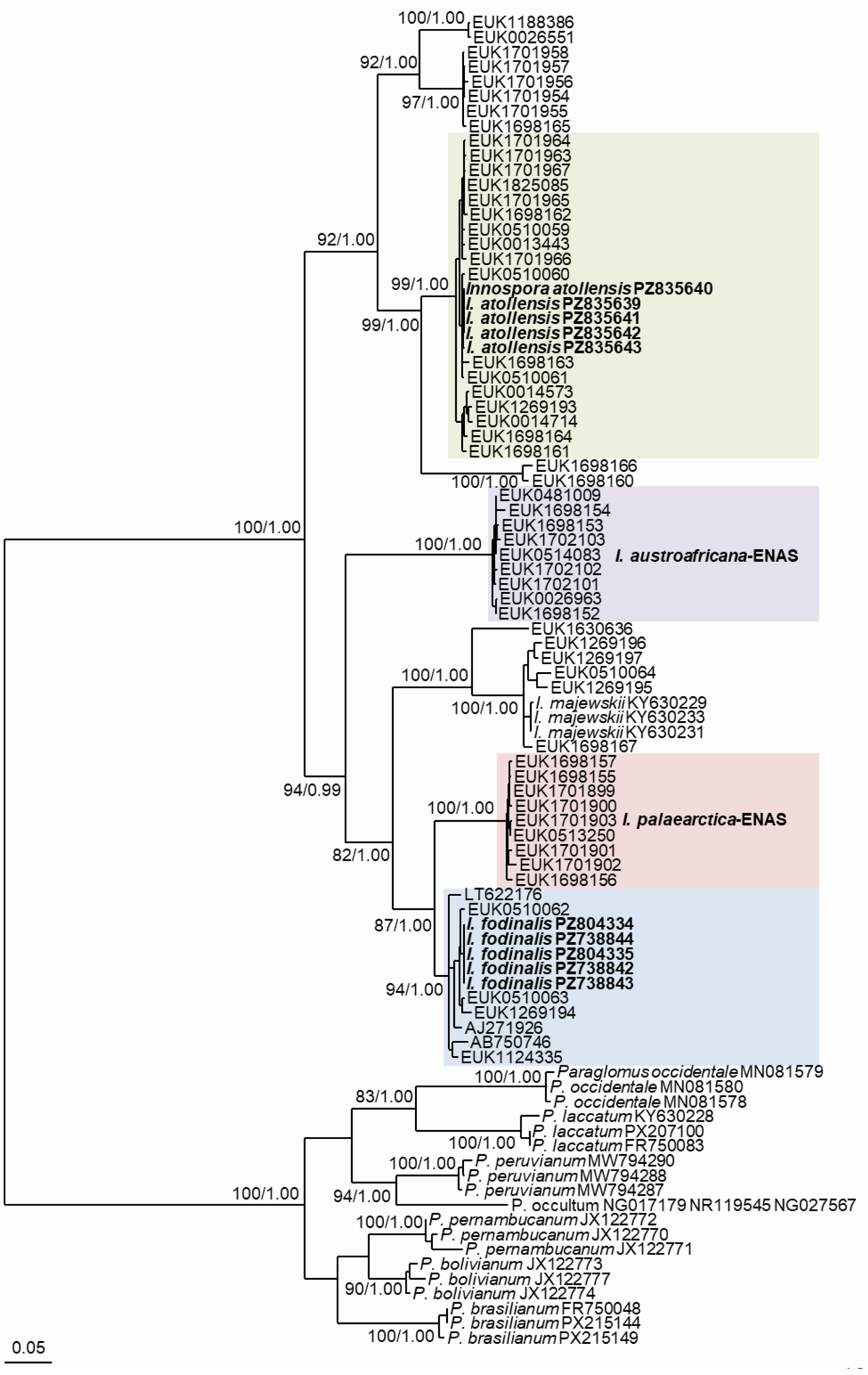
Phylogenetic tree showing the topology recovered from maximum-likelihood and Bayesian inference analyses of a dataset comprising *Innospora* species, 58 environmental sequences retrieved from EUKARYOME and GenBank, and *Paraglomus* species as the outgroup. For the Bayesian inference, all convergence diagnostics indicated that convergence had been achieved (ASDSF = 0.009; average PSRF for parameter values = 1.001). Sequences of two newly described *Innospora* species are indicated in bold. Coloured boxes are used to delimit the species clades, including environmental sequences, and the environmental lineages designated as DFT. Values on the branches represent maximum-likelihood bootstrap support (BS) and Bayesian posterior probabilities (PP); only BS ≥ 80% and PP ≥ 0.97 are shown. The scale bar indicated 0.05 substitutions per site.

Two fully supported environmental lineages met the criteria for classification according to the International Standard to Catalogue Dark Fungal Taxa (ISCDFT), such as the presence in at least three independent samples. One lineage (% dissimilarity among its sequences = ca. 1.9%), hereafter referred to as *I. austroafricana*-ENAS, was recorded only at South African sites, whereas the other lineage (% dissimilarity among its sequences = ca. 3.3%), hereafter referred to as *I. palaearctica*-ENAS, showed an Euro-Mediterranean distribution. The phylogeny, including environmental lineages, revealed several additional well-supported relationships among *Innospora* lineages, and broader clades. *Innospora fodinalis* and *I. palaearctica*-ENAS engaged in a supported (ML = 87, BI = 1.00) sister relationship with a 45S sequence dissimilarity of ca. 9.7%. Together with *I. majewski*i and *I. austroafricana*-ENAS, they formed a supported group (ML = 94, BI = 0.99) distinct from a clade (ML = 92, BI = 1.00) populated by *I. atollensis* and a few other environmental lineages. Alignments and tree files for Fig. 2 and Fig. 3 are provided as Suppl. material 2. Sequence accession numbers with references are provided as Suppl. material 3.

### Description of new species

#### Innospora atollensis

T. Crossay, M. Wong, sp. nov. Figure 4A–F.

**Figure 4.**
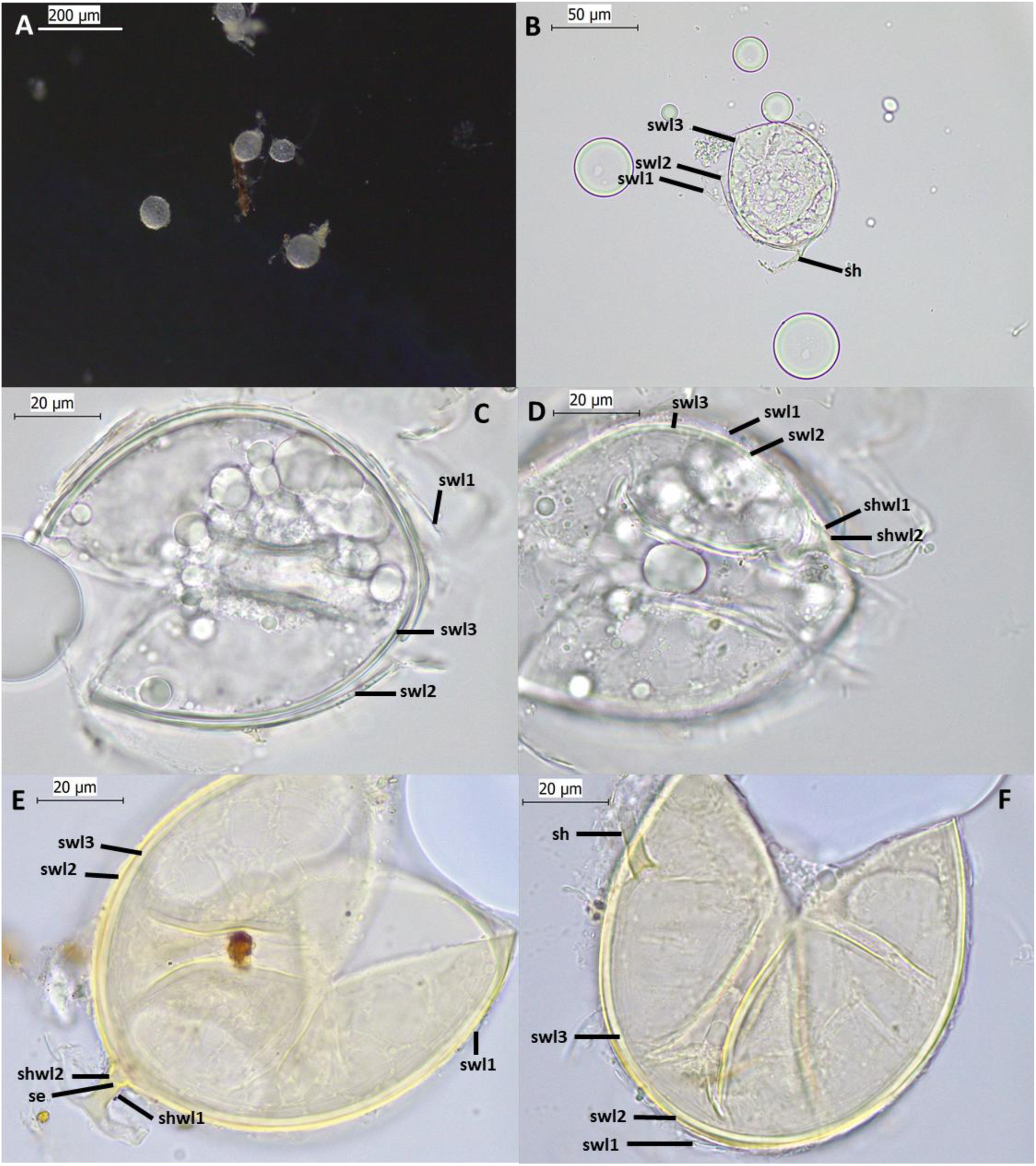
*Innospora atollensis.* **(A)** Intact spore and **(B, C, D, E, F)** Crushed spores in PVLG **(B, C, D)** and PVLG + Melzer **(E, F)**. **(B–F)** Spore wall layers (swl) 1–3. **(B, D, E)** Subtending hypha (sh) with two wall layers (shwl) 1–2, and septum (se). Scale bars:**(A)** = 200 μm, **(B)** = 50 μm, **(C–F)** = 20 μm.

#### MycoBank No

MB number will be added after review.

#### Etymology

Latin, *atollensis*, referring to an organism that lives on an atoll.

#### Specimens examined

Isolated from rhizospheric soil of a greenhouse pot of a single– species culture propagated on *Sorghum vulgare* at the laboratory Aura Pacifica in New Caledonia, Noumea, June 2024, T. Crossay. This single–species culture was originally inoculated with 100 spores isolated from the rhizospheric soil of *Ficus carica* sampled at a naturally vegetated site on Rangiroa Island (14°58’24.3”S; 147°37’28.4”W) in French Polynesia, in October 2022, T. Crossay. Holotype deposited at the Mycological Herbarium of the National Museum (France, Paris) in MNHN–PC–will be added after review. Related environmental sequences include: EUK0013443, EUK0014573, EUK0014714, EUK0510059, EUK0510060, EUK0510061, EUK1269193, EUK1698161, EUK1698162, EUK1698163, EUK1698164, EUK1701963, EUK1701964, EUK1701965, EUK1701966, EUK1701967, EUK1825085.

#### Diagnosis

It differs from the other species of the genus *Innospora* in (i) the presence of a close septum, (ii) the thickness of spore wall layer 3, and (iii) the nucleotide composition of 45S sequences. Separation of *Innospora atollensis* from other *Innospora* species based on the ITS region (ITS2 positions 182–202 TTTTAGATAGGGTGTGGCAA; no mismatch allowed) and LSU (positions 564–587 TAGACCGGAGCAGATGCAATTGAC; no mismatch allowed) (Suppl. material 4A).

#### Description

Glomerocarps unknown. Glomerospores produced singly in soil, blastically at the tip of a sporogenous hypha. Spores globose with one subtending hypha, hyaline, (45–)60(–65) μm diam. (Figure 4A). Spore wall composed of three layers (Figure 4B-F). Layer 1 (SWL1), evanescent, usually slightly roughened on its upper surface, hyaline, (0.5–)0.8(–1.1) μm thick (Figure 4B-F). Layer 2 (SWL2) uniform, smooth, permanent, laminated, light yellow (1A2) (Figure 4B-D), (1.2–)1.5(– 2.4) μm thick. Layer 3 (SWL3) uniform (without visible sublayers), flexible, smooth, hyaline (Figure 4D, E), (1.9–)2.4(–2.9) μm thick, generally not adherent to layer 2. None of the spore wall layers stains in Melzer’s reagent (Figure 4E, F). Subtending hypha hyaline, straight or curved, cylindrical; (2.9–)3.5(–3.9) μm wide at spore base (Figure 4D-F). Wall of subtending hypha hyaline ; (1.2–)1.5(–1.7) μm thick at spore base; continuous with layers 1 and 2 of spore wall (Figure 4E). Pore (1.5–)2(–2.4) μm in diameter, occluded by a plug formed by layer 2 at the spore base (Figure 4E). Spore content of hyaline oily substance (Figure 4B-D). Germination unknown. Forming typical mycorrhiza with arbuscules, coils and intraradical hyphae staining faintly with Trypan Blue in cortical roots of *Sorghum vulgare*.

#### Distribution and habitat

In the field, *I. atollensis* has only been reported from the type locality on Rangiroa Island, French Polynesia. However, environmental sequences recorded in the EUKARYOME v.2.1 database suggested a broader distribution, with records from Cuba, the United States, Australia, and Guadeloupe (Suppl. material 1). These sequences originate from diverse biomes, including subtropical woodland, subtropical broadleaf forest, tropical broadleaf forest, and urban environments.

#### Innospora fodinalis

F. Magurno, S. Szczecinska, D. Di Loreto, T. Crossay, sp. nov. Figure 5 A–F.

**Figure 5.**
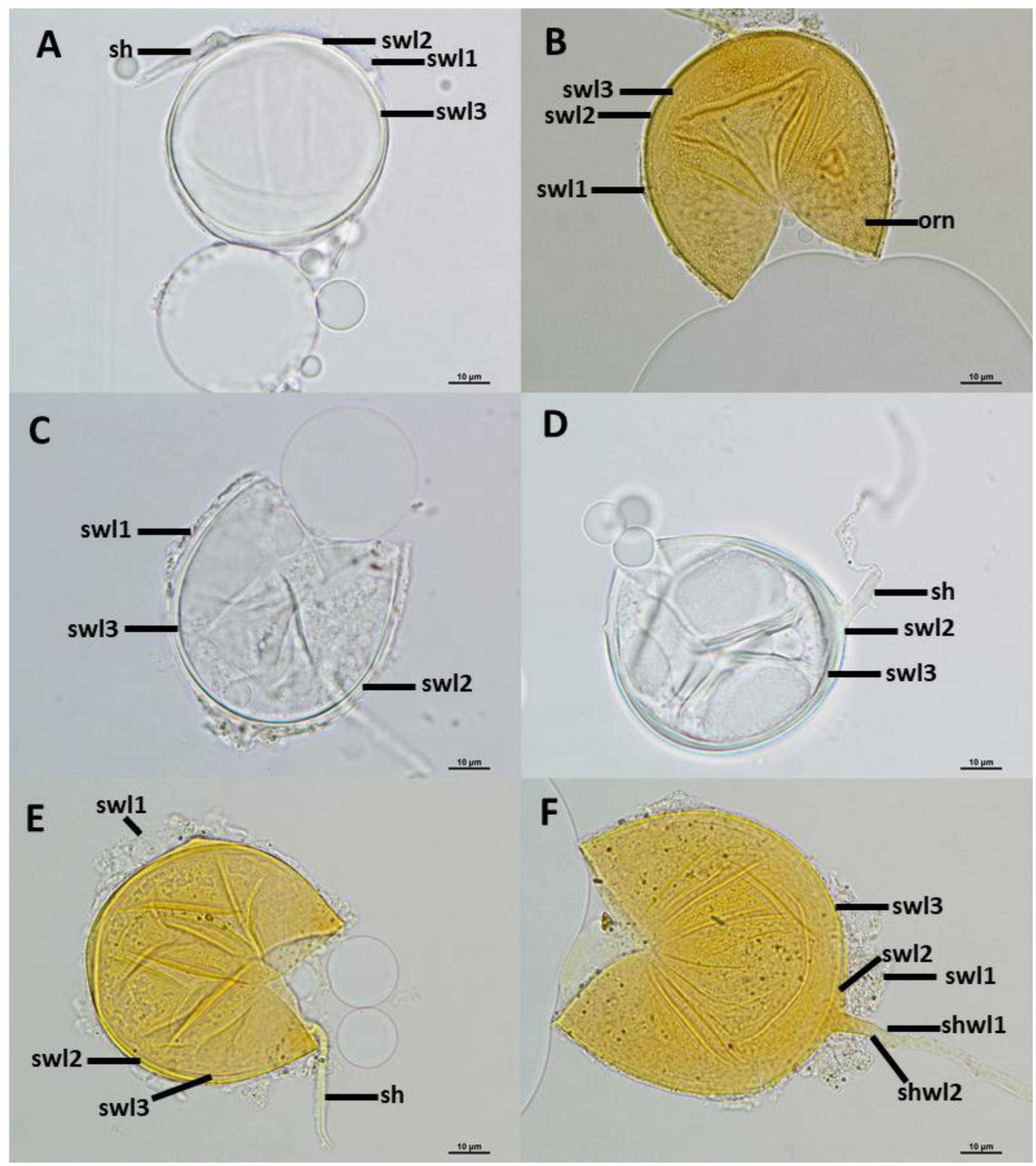
*Innospora fodinalis.* **(A)** Intact spore and **(B, C, D, E, F)** Crushed spores in PVLG **(A, C, D)** and PVLG + Melzer **(B, E, F)**. **(B)** ornamentation (orn) and **(A–F)** spore wall layers (swl) 1–3. **(A, D, E, F)** Subtending hypha (sh) with two wall layers (shwl) 1–2. Scale bars:**(A–F)** = 10 μm.

#### MycoBank No

MB number will be added after review.

#### Etymology

from the Latin *fodina* (= mine), referring to the occurrence of the species in post mining sites.

#### Specimens examined

isolated from the substrate of a single species culture with *Plantago lanceolata* as host plant. This single–species culture was originally inoculated with 1 spore isolated from a trap culture set up with rhizospheric soil of *Tanacetum vulgare*. The soil sample was collected by Franco Magurno at the post mining coal spoil heap of Zabrze Sosnica Makoszowy (50°16’21.5”N; 18°44’42.7”E), in the proximity of Zabrze, Poland, October 2019. Holotype deposited at the KTU Herbarium of the University of Silesia in Katowice (Poland), Slide No. 0166579. Related environmental sequences include: AB750746, AJ271926, EUK0510062, EUK0510063, EUK1124335, EUK1269194, LT622176.

#### Diagnosis

It differs from the other species of the genus *Innospora* in (i) the presence of granular ornamentation on spore wall layer 1 and (ii) the nucleotide composition of 45S sequences. Separation of *Innospora fodinalis* from other *Innospora* species based on the ITS region (ITS2 positions 12–28 AATACTACGAAAAATGT; no mismatch allowed) and LSU (positions 459–482 GAGCTTTTTCGCGAAAGCTTGTGA; no mismatch allowed) (Suppl. material 4B).

#### Description

Glomerocarps unknown. Glomerospores produced singly in soil, blastically at the tip of a sporogenous hypha. Spores globose with one subtending hypha, hyaline, (48–)51(–61) μm diam. (Figure 5A). Spore wall composed of three layers (Figure 5A-F). Layer 1 (SWL1), evanescent, ornamented with small granular excrescences (0.3–)0.5(–0.7) μm diam., hyaline, (1.1–)1.6(–2.4) μm thick (Figure 5B, C, E, F). Layer 2 (SWL2) uniform, smooth, permanent, hyaline (Figure 5A-F), (0.7–)0.8(–0.9) μm thick. Layer 3 (SWL3) uniform (without visible sublayers), flexible, smooth, hyaline (Figure 5A-F), (0.9–)1.1(–1.5) μm thick, generally adherent to layer 2. None of the spore wall layers stains in Melzer’s reagent (Figure 5B, E, F). Subtending hypha hyaline, straight or curved, cylindrical; (1.9–)2.1(–2.6) μm wide at spore base (Figure 5D-F). Wall of subtending hypha hyaline ; (1.0–)1.4(–1.6) μm thick at spore base; continuous with layers 1 and 2 of spore wall (Figure 5F). Pore (0.8–)0.9(–1.1) μm in diameter, open, formed by layers 2 at the spore base (Figure 5D, F). Germination unknown. Forming typical mycorrhiza with arbuscules, coils and intraradical hyphae staining strongly with ink-vinegar solution in cortical roots of *Plantago lanceolata*.

#### Distribution and habitat

In the field, *I. fodinalis* is represented by an isolate obtained from a post-coal mining wasteland in Poland. Additional environmental sequences from EUKARYOME v.2.1 and GenBank expand the occurrence of the species to Poland, Japan, Estonia, China, Russia, and France (Suppl. material 1). The recorded habitats include anthropogenically disturbed environments, such as post-mining sites and urban areas, temperate grasslands and sand dunes.

### Designation of new DFT

#### Innospora palaearctica-ENAS F

Magurno, K. Esmaeilzadeh-Salestani, L. Tedersoo, sp. nov.

#### Identifier

MBDFT will be added after review.

#### UNITE DFT-SH DOI

will be added after review.

#### Etymology

From *palaearcticus* (of the Palaearctic realm), referring to its distribution across Europe, North Africa and the Caucasus.

#### Diagnosis

Differs from the sister species *Innospora fodinalis* and from other species of *Innospora* based on the ITS2 subregion (positions 61–80: CCGAATAATAGTAATTATTA; one mismatch allowed) and LSU (positions 459–480: GAGGTTTTTCGAAAGCTCGTGA; no mismatch allowed) (Suppl. material 4C).

#### Reference sequence (legitype)

EUK1698156 = INSD-accession. Morocco, Al-Hoceima, 35°11’00”N, 4°08’38”W, 454 m a.s.l. (GSMc plot G4048, vouchered sample TUE000622, DNA sample TUE100622), Mediterranean forest biome, soil, 2014-12-09, leg. Leho Tedersoo.

#### Additional sequences

EUK1701899, EUK1698155, EUK1698157, EUK1701902, EUK0513250, EUK1701901, EUK1701903, EUK1701900.

#### Ecology & distribution

Found in the soil of temperate coniferous, temperate broadleaf and Mediterranean forests, temperate shrubland, and montane grassland in Croatia, Estonia, Georgia, Italy, and Morocco (Suppl. material 1), between ca. 31°N to 58°N and from 15 to 3158 m a.s.l.

#### Notes

Forms a distinct, well-supported lineage within *Innospora* (96.6% mean intraspecific identity). Its closest relative is *I. fodinalis*, with 90.3% identity. The nine sequences represent six independent samples from six localities.

#### Innospora austroafricana-ENAS

F. Magurno, K. Esmaeilzadeh-Salestani, L. Tedersoo, sp. nov.

#### Identifier

MBDFT will be added after review.

#### UNITE DFT-SH DOI

will be added after review.

#### Etymology

From *austro-*(southern) and *africanus* (African), referring to its occurrence in South Africa.

#### Diagnosis

Differs from other species of *Innospora* based on the ITS2 subregion (positions 260–279: AAAAAAGATCGAAAAAGATC; no mismatch allowed) and LSU (positions 122–143: TGGCTGCAATGAGCAGTCCGAC; no mismatch allowed) (Suppl. material 4D).

#### Reference sequence (legitype)

EUK1698154 = INSD-accession. South Africa, St. Lucia, 8°23’02.8”S, 32°24’33.5”E, 23 m a.s.l. subtropical broadleaf forest biome, soil, 2016-03-12, leg. Sten Anslan (GSMc plot S655, vouchered sample TUE003159, DNA sample TUE103159).

#### Additional sequences

EUK1698152, EUK1702102, EUK1698153, EUK0481009, EUK0514083, EUK0026963, EUK1702101, EUK1702103.

#### Ecology & distribution

Found only in South Africa, in the soil of subtropical broadleaf forests, at four coastal localities (Platbos forest, Wilderness, St. Lucia, Cape Vidal; Suppl. material 1). Ranges from approximately 28°S to 35°S and from 23 to 227 m a.s.l.

#### Notes

Forms a highly cohesive, well-supported lineage within *Innospora* (98.1% mean intraspecific identity). It is strongly separated from all other sequences: the nearest sequences outside the clade share at most 83.2% identity, and none of them is assigned to a named species. The nine sequences represent four independent samples, six of them from the Platbos forest.

## Discussion

The description of the four *Innospora* species, *I. atollensis* and *I. fodinalis*, as well as *I. austroafricana*-ENAS and I. *palaearctica*-ENAS, expanded our knowledge of *Innospora*, a monospecific genus in the *Paraglomeraceae* represented so far only by *I. majewskii*. The newly described species and dark fungal taxa also exhibited distinct biogeographic patterns. *Innospora atollensis* showed a tropical–subtropical trans-Pacific distribution extending into the Caribbean, with records from French Polynesia, Australia, Cuba, Guadeloupe and the United States.

Its occurrence on a coral atoll implies long-distance dispersal, because atoll islets are low-lying land built from coral debris and sand (Duvat et al. 2020; Montaggioni et al. 2023). This dispersal may have been natural or associated with the introduction of plants such as *Ficus carica*, the host plant at the sampling site. *Innospora fodinalis* was similarly broadly distributed across Eurasia, whereas *I. palaearctica*-ENAS was restricted to the Euro-Mediterranean region and *I. austroafricana*-ENAS to coastal subtropical forests of South Africa. Although *I. fodinalis* and its sister lineage *I. palaearctica*-ENAS partially overlap geographically, they appear to occupy different habitats. *Innospora palaearctica*-ENAS occurred mainly in forest and shrubland soils, whereas *I. fodinalis* was found mostly in open, disturbed habitats. About half of the occurrences of *I. fodinalis* came from post-mining sites: besides the type locality, these were the former lead and zinc mining site of Les Avinières in France (Sánchez-Castro et al. 2017) and a 20-year-old zinc waste site in southern Poland (Turnau et al. 2001). Such substrates are coarse, nutrient-poor, acidic and saline, and often contaminated with heavy metals (Woźniak 2010). The detection of *I. fodinalis* at such sites suggests that the species may have traits enabling it to tolerate these conditions. These patterns should nevertheless be interpreted with caution. Environmental records come from a few long-read datasets with uneven geographic coverage. The ranges reported here are therefore minimum estimates. Nevertheless, the detection of *I. austroafricana*-ENAS at four well-separated areas of South Africa, and its absence from the thousands of sites sampled worldwide in EUKARYOME, suggests that this lineage may be endemic to the region.

Similarly to *I. majewskii*, the two isolates representing the two new species showed a very low intra-isolate diversity (<0.5%). However, due to the lack of a chromosome-scale genome assembly, such as those available for *Rhizophagus irregularis* (Yildirir et al. 2022) and *Gigaspora margarita* (Mugambi et al. 2026), it was not possible to verify the number and exact divergence of paralog copies. However, when the analysis was expanded to include environmental sequences, maximum 45S sequence dissimilarity within clades recognized as species reached approximately 4%. Within their respective species-level clades, sequences from each of the two isolates nevertheless formed distinct, well-supported subclades. Although in *I. atollensis* several additional well-supported environmental subclades were recovered within the broader species-level clade, the available data were insufficient to determine whether any of these lineages should be recognized as separate species.

In contrast to the extremely low intra-isolate diversity observed within *Innospora* isolates, interspecific sequence divergence within the genus was remarkably high, reaching levels comparable to those typically observed between families, or even between orders. However, even if the phylogeny including environmental sequences revealed several well-supported relationships among *Innospora* lineages, we decided not to erect new genera. According to the isolate-based phylogeny, the three *Innospora* species would have to be placed in three distinct genera. Taking the environmental lineages into account, such a separation would imply the recognition of at least four genera, three of them monospecific (*I. majewskii*, *I. atollensis*, *I. austroafricana*-ENAS). Furthermore, the high sequence dissimilarity among *Innospora* species appears to be strongly “inflated” by the ITS1 and ITS2 subunits of rDNA sequences. Indeed, when sequence comparisons were based on LSU or RPB1, divergence among *Innospora* species was comparable to, or lower than, that observed among *Paraglomus* species. For these reasons, fragmentation of *Innospora* into multiple genera would appear premature and poorly justified by the currently available molecular evidence.

Spores of the newly described species (*I. atollensis*, *I. fodinalis*) and of *I. majewskii* are characterized by morphological traits shared with species of other genera producing glomoid spores (Błaszkowski et al. 2017). Nevertheless, a few morphological characteristics allow them to be distinguished. For an experienced observer, examination under a stereomicroscope can narrow down the identification to the genera *Paraglomus*, *Pervetustus*, *Innospora*, or *Archaeospora*, based on the lack of spore pigmentation (except for *P. bolivianum* producing yellow brown to brown spores), their small size, their production singly in soil, and their tendency to float in water (Morton and Redecker 2001; Błaszkowski et al. 2012; de Mello et al. 2013). Preparation of spores on microscope slides under a coverslip followed by careful microscopic examination is then necessary to classify these species within the genus *Innospora*. First, the formation of one-walled spores clearly distinguishes *Innospora* spp. from *Paraglomus* spp., which produce two-walled spores (Błaszkowski et al. 2017). *Innospora* spores can also be distinguished from those of *Pervetustus simplex*, which has two layers (vs three layers for spores of *Innospora* spp.), and a spore wall 1.4 to 1.9 times thicker (means of spores wall thickness : *Pe. simplex* 6.6 μm, *I. majewskii* 4.7 μm, *I. fodinalis* 3.5 μm, *I. atollensis* 4.7 μm; Błaszkowski et al. 2012, 2017). As a consequence, spores of *Pe. simplex* are less fragile and more flexible than those of *Innospora* spp., and thus less likely to break or open when mounted under a coverslip (Crossay, pers. observ.). Spore wall thickness comparisons reported above highlight a likely inversion error in Błaszkowski et al. (2017; p. 408, line 14). Finally, *Innospora* spp. spores can be distinguished from those of *Archaeospora* spp. by their mode of formation. *Innospora* spp. typically produces glomoid spores at the tip of a subtending hypha, as characteristic of *Paraglomerales*, whereas the acaulosporoid spores of *Archaeospora* spp. develop laterally on, or within, the hypha subtending a sporiferous saccule (Sieverding and Oehl 2006).

Morphological differentiation among the three *Innospora* species is more difficult; *Innospora fodinalis* is the most readily recognizable because the first layer of its spore wall harbors small granular excrescences on its surface (Fig. 5B) which are absent in the other two species (Błaszkowski et al. 2012; Figs. 4 and 5). The spores of *I. atollensis* differ from those of *I. majewskii* and *I. fodinalis* in two morphological characteristics. The first and most evident is the thickness of the third wall layer, which is on average 2.4 times thicker than that of *I. majewskii* and 1.6 times thicker than that of *I. fodinalis* (Błaszkowski et al. 2012; Figs. 4 and 5). The second characteristic is the presence, at the spore base, of a pore occluded by a plug formed by wall layer 2 (Fig. 4E); this structure has not been observed in the other two species, both of which having an open pore at the spore base (Błaszkowski et al. 2012; Fig. 5).

## Conclusions

The present study substantially expands the known diversity of the monospecific genus *Innospora*, with the description of two new species and two dark fungal taxa. The genus is characterized by a striking contrast between the apparently very low intra-isolate variability and high interspecific divergence, largely concentrated in the ITS-rDNA regions. Consistent with the generally limited morphological differentiation among members of *Paraglomerales*, *Innospora* species are distinguished mainly by molecular divergence and phylogenetic inference, while morphological differences remain subtle.

## Author contributions

FM: Conceptualization, Data curation, Formal analysis, Investigation, Methodology, Supervision, Writing – original draft, Writing – review & editing; SS: Data curation, Formal analysis, Investigation, Methodology, Writing – review & editing; DD: Data curation, Formal analysis, Investigation; DF: Formal analysis, Writing – original draft, Writing – review & editing; KE-S: Formal analysis, Writing – original draft, Writing – review & editing; MW: Writing – review & editing; MF: Visualization; LT: Conceptualization, Funding acquisition, Supervision, Writing – review & editing; TC: Conceptualization, Data curation, Formal analysis, Funding acquisition, Investigation, Methodology, Supervision, Writing – original draft, Writing – review & editing.

## Acknowledgements

We respectfully acknowledge the late Christine Wong (DAG) for her contributions to this project. We equally thank Elena Gorchakova (UICN) and Laurent Maunas (DAG) for their contributions.

